# Degraded neural coding of temporal fine structure with age predicts effortful listening in multi-talker environments

**DOI:** 10.1101/2025.11.17.688858

**Authors:** Leslie Zhen, Satyabrata Parida, Jacie R. McHaney, Maggie E. Zink, Bharath Chandrasekaran, Aravindakshan Parthasarathy

## Abstract

Middle age represents a critical window for early detection of neurophysiological decline. Hearing loss is increasingly recognized as both an early marker of neural degeneration and a modifiable risk factor for dementia. Yet many adults report difficulty understanding speech in noise despite normal audiograms, highlighting the limitations of current clinical tests that fail to capture the underlying physiology or effort required for real-world listening. Beyond hearing thresholds, speech comprehension in complex environments depends on precise neural encoding of temporal fine structure (TFS) cues that convey pitch and spatial information. Here, we use a noninvasive EEG-based measure of neural phase-locking (frequency modulation following responses or FMFRs) to quantify TFS encoding in young and middle-aged adults with normal hearing thresholds. Middle-aged listeners exhibited reduced FMFR amplitudes and shallower discriminability slopes, reflecting diminished neural synchrony despite preserved hearing thresholds. Using a multi-talker speech task we further found that pupil-indexed listening effort was significantly greater in middle-aged adults despite matched accuracy across groups. Further, increases in listening effort were predicted by decreases in TFS encoding. Together, these results reveal that degraded neural encoding of TFS underlies subclinical listening difficulties and increased cognitive load, establishing the FMFR as a sensitive biomarker of hidden auditory neural decline.

**Significance:** Understanding speech in noisy environments depends on precise neural encoding of temporal fine structure (TFS) cues. Using a noninvasive EEG metric, the FMFR, we show that neural TFS coding in the peripheral auditory system declines markedly by midlife, even when hearing thresholds and speech performance are normal. These neural deficits predict elevated pupil-indexed listening effort during multi-talker speech perception, revealing that subclinical degradation of temporal coding increases cognitive load in everyday listening. The FMFR thus provides a promising biomarker for early auditory neural decline and its downstream cognitive consequences.

## Introduction

Middle age marks a critical window for detecting the earliest signs of sensory and neural decline (Dohm-Hansen et al., 2024). Hearing loss, long regarded as an inevitable consequence of aging, is now recognized as a modifiable risk factor for dementia and a key indicator of broader neurodegenerative vulnerability (Livingston et al., 2017, 2020). Yet a persistent clinical paradox remains - many middle-aged adults report difficulty understanding speech in noise despite having normal audiometric thresholds (Hind et al., 2011; Spehar and Lichtenhan, 2018; Parthasarathy et al., 2020; Cancel et al., 2023).

Accurate speech perception in multi-talker conditions relies on the precise neural encoding of rapid temporal fine structure (TFS), which provides pitch and spatial information critical for understanding speech in noise and reverberation (Moore, 2008; Borjigin and Bharadwaj, 2025). Deficits in sensitivity to TFS cues are implicated in speech perception in noise (SPIN) difficulties, especially among listeners with hearing loss and cochlear implant users, the latter notably lacking TFS representation entirely (Shannon et al., 1995; Lorenzi et al., 2006, 2009; Moore, 2008; Strelcyk and Dau, 2009). Additionally, even among normal-hearing adults, individual variability in TFS processing predicts speech-in-noise performance deficits, underscoring the need for objective, sensitive assessments of neural TFS encoding (Ruggles et al., 2012; Parthasarathy et al., 2020). Behavioral assays of TFS sensitivity, such as frequency-modulation (FM) detection, can be contaminated by recovered envelope cues created by cochlear filtering, obscuring the contribution of true phase-locked coding (Verschooten et al., 2019). To circumvent these methodological limitations, we recently validated frequency modulation following responses (FMFRs), an electroencephalography (EEG)-based direct measure of neural phase-locking to low-frequency FM stimuli, precisely capturing TFS encoding fidelity, free from confounding envelope cues. Individual neural differences in TFS processing measured using FMFRs predict multi-talker speech intelligibility in normal-hearing young adults (Parthasarathy et al., 2020).

Moreover, behavioral accuracy alone often underestimates perceptual strain, as listeners can maintain performance at the cost of greater cognitive effort (McHaney et al., 2024; Zink et al., 2024). Task-evoked pupillary dilations provides a sensitive index of this effort, reflecting the engagement of central arousal systems during demanding listening (Kahneman and Beatty, 1966; Pichora-Fuller et al., 2016). Task-evoked changes in pupil diameter can hence provide a sensitive metric of overall cognitive load that reflects many critical elements of speech perception in noise, including task complexity, difficulty, memory load, selective attention, acoustic degradation, and sentence recognition, integration and repair (Zekveld et al., 2011; Kramer et al., 2013; Zekveld and Kramer, 2014; Winn et al., 2015; Winn, 2016; Parthasarathy et al., 2020; McHaney et al., 2024; Zink et al., 2024).

Here we unite these domains of temporal coding fidelity and listening effort within a single experimental framework. Using the FMFR as an electrophysiological measure of neural TFS encoding, we quantify the fidelity of TFS encoding in young and middle-aged adults with clinically normal hearing. We pair this with pupillometric measures of listening effort during multi-talker speech comprehension with matched behavioral outcomes across the age groups, dissociating listening effort from behavioral performance. We hence test whether degraded neural encoding predicts increased cognitive load despite preserved behavioral performance. This integrative approach bridges the gap between peripheral neural coding and listening effort, providing a mechanistic account of how subclinical auditory decline manifests as listening difficulties in everyday communication.

## Materials and Methods

### Participants

#### Recruitment

Eighty-three adults (26 male; age range: 18–55 years) were recruited through the Pitt+Me Research Registry, the University of Pittsburgh Department of Communication Science and Disorders research participant pool, and community advertisements. Participants were compensated for time and travel and received an additional incentive upon completion of all sessions. All procedures were approved by the University of Pittsburgh Institutional Review Board (IRB #21040125), and informed consent was obtained from all participants.

#### Otoscopy

Ear canals and tympanic membranes were examined with a Welch Allyn otoscope. Participants with excessive cerumen, drainage, or other conductive abnormalities were excluded. *Audiometry*. Air-conduction thresholds were obtained using a MADSEN Astera^2^ audiometer (Otometrics/Natus Medical, Middleton, WI) with foam insert ear tips. The modified Hughson– Westlake procedure was used, and tones were presented at octave frequencies from 0.25 to 8 kHz. Extended high-frequency thresholds (8, 12.5, and 16 kHz) were measured using Sennheiser HDA 300 circumaural headphones. Due to audiometer limitations, thresholds greater than 35 dBHL at 16kHz were marked as 35dBHL.

#### Loudness Discomfort Levels

Binaural LDLs were measured using warble tones (0.5–3 kHz) delivered via insert earphones. Participants rated loudness on a 1–7 scale, with “7” indicating intolerable loudness.

#### Distortion Product Otoacoustic Emissions (DPOAEs)

Outer hair cell function was assessed using an ER-10D DPOAE probe (Etymotic Research, Elk Grove, IL). Primary tones (L_1_ = 75 dB SPL, L_2_ = 65 dB SPL) were presented from 500 Hz–16 kHz in 8 blocks of 24 alternating-polarity sweeps per ear. *Noise Exposure History*. Self-reported annual noise exposure was estimated using the Noise Exposure Questionnaire (NEQ;). Cumulative exposure was expressed as LAeq8760h, representing the annual equivalent continuous sound level.

#### Inclusion criteria

Participants had normal cognition (Montreal Cognitive Assessment, MoCA ≥ 25; Nasreddine et al., 2005), clinically normal hearing thresholds (≤25 dB HL, 250–8000 Hz), and no severe tinnitus (Tinnitus Handicap Inventory < 38). Loudness discomfort levels (LDLs) exceeded 80 dB HL at 0.5, 1, and 3 kHz. Participants were native or fluent speakers of American English. Two age-defined cohorts were invited for full testing: 36 young adults (YA, 18–25 years; Mean age 21.1+ 1.8yrs, 10 male) and 36 middle-aged adults (MA, 40–55 years; Mean age 46.5+ 4.7 years, 11 male).

### Frequency-Modulation Following Responses (FMFRs)

#### Recording setup

EEG recordings were obtained in an electrically shielded, sound-attenuating chamber. Participants reclined comfortably and were instructed to minimize movement; arousal state was monitored but not constrained. Recording sessions lasted approximately 3 h, with breaks as needed. Neural activity was collected using a 64-channel BioSemi ActiveTwo system at 16 kHz sampling rate. Two foil tiptrodes (Etymotic) were placed in the ear canals, and six additional cup electrodes were positioned at horizontal and vertical EOG sites and both earlobes.

Signals were acquired with a common-average reference. Stimuli were delivered through calibrated ER-3C insert earphones (Etymotic) using a TDT real-time processor (sampling rate = 100 kHz), synchronized with BioSemi acquisition at 25 kHz under custom LabVIEW control.

#### Stimuli and paradigm

The FMFR was elicited by 1-s sinusoidal frequency-modulated (FM) tones with a 520-Hz carrier, 2-Hz modulation rate, and deviation depths of 0 (pure-tone), 2, 5, 8, and 10 Hz. Each stimulus had 5-ms raised-cosine onset and offset ramps and was presented every 1.19 s at 85 dB SPL. Polarity alternated across trials, with 200 epochs collected per polarity.

#### Neural signal extraction

Analyses focused on the Fz–tiptrode montage, which optimally captures low-frequency phase-locked responses originating in the auditory nerve and brainstem(McHaney et al., 2024). Because cochlear neural responses to low-frequency tones are phase-sensitive and invert with stimulus polarity, the summed response to alternating polarities is effectively rectified and periodic at twice the carrier frequency (Lichtenhan et al., 2014). FMFR magnitude was therefore quantified at 2 × carrier frequency.

#### FMFR power and neural discriminability metrics

To quantify how well the neural response (FMFR) tracked the frequency-modulation trajectory, we employed spectrally specific analyses, which yield optimal spectral resolution for dynamic signals with known trajectories (Parida et al., 2021, 2024). Briefly, the FMFR was first band-pass filtered between 300 and 2600 Hz. Next, the filtered signal was frequency demodulated along the desired FM trajectory, which projects the desired signal onto 0 Hz. Here, we estimated the power along the frequency trajectory that is two-times the carrier frequency in the summed-FFR response. Next, we estimated the multi-taper spectrum (with time-bandwidth half product of 1.5 using the Matlab function *pmtm*) and computed the power near 0 Hz (6-Hz bandwidth). We estimated noise floor as the power that is + 15 Hz of the demodulated power trajectory near 0 Hz. We estimated this FM power for each FM deviation depth. The neural discriminability index was computed as the relative power (in dB) for a given FM deviation depth relative to the pure sinusoidal condition (where the FM trajectory is constant).

### Speech perception in noise

#### Digits Comprehension Task

The task was divided into two sections of trials: 1) practice and 2) test. In the practice section, participants completed 8 practice trials. The first two practice trials were introductory trials to the target male speaker (F0 = 115 Hz) as he produced a string of four randomly selected monosyllabic digits (digits 1–9, excluding the bisyllabic ‘7’) with 0.68 s between the onset of each digit. After these two introductory trials, participants then completed six trials of the target speaker masked with two additional speakers (male, F0 = 90 Hz; female, F0 – 175 Hz) that produced randomly selected digits with matched target-onset times. The two competing speakers could not produce the same digit as the target speaker or each other, but otherwise digits were presented at random. The masker stimuli were presented at 60 dB SPL. The intensity of the target speaker was modulated to obtain the desired signal-to-noise ratio level. During the practice trials, there were two trials at each SNR of 20 dB, 16 dB, and 9 dB in descending order. At the start of each trial, a 1000 Hz beep was presented to signal the start of the trial, followed by two seconds of silence, then the digit stimuli, two seconds of silence after the fourth digit, and a final 1000 Hz beep to signal the end of the trial. At the presentation of the beep at the end of the trial, participants were prompted to verbally repeat the target 4-digit sequence, which was recorded by the experimenter.

Upon completion of the practice section, the participants moved onto the test section. The test section consisted of 8 blocks with 5 trials per block. Within a block, all trials were presented at the same SNR level of 9, 6, 3, or 0 dB. The test trials followed the same timing procedure as the practice trials, including the beeps to signal the start and end of each trial. Two blocks were presented per SNR level, and blocks were presented in random order. Participants received a self-timed break at the end of each trial. Stimulus presentation was controlled using MATLAB 2018b (MathWorks, Inc.). Trials were scored correct if all four digits repeated by the participant matched the target speaker sequence in the correct order. The behavioral response curve was constructed using percent correct as a function of SNR. Stimuli were presented through calibrated Sennheiser circumaural headphones.

### Pupillometry

#### Acquisition

Monocular left eye pupillary responses were recorded during the digits comprehension task using an Eyelink 1000 Plus Desktop Mount (SR Research) with a chin and forehead rest for stabilization. Data were recorded in arbitrary units at a sampling rate of 1000 Hz. Luminance of the visual field was controlled by setting consistent room lighting across all participants. Nine-point eye tracker calibration was performed prior to the start of the experiment. We included a mandatory fixation criterion at the start of each trial to help control for the effects of saccades, which affect pupillary measurements, and to minimize pupil foreshortening errors (Hayes and Petrov, 2016). Participants were required to fixate on a cross on the screen for a minimum of 500 ms to initiate the start of each trial. Manual drift correction was performed by the experimenter at the end of each trial to ensure high-quality tracking of the pupil throughout the digits comprehension task and to account for possible shifts in position as a result of providing a verbal response on each trial.

#### Preprocessing

Raw pupillary data recorded while participants listened to digits sentences were processed in R (R Core Team, 2022). Data were first downsampled to 50 Hz, then processed to remove noise from blinks and saccades. Any trial with more than fifteen percent of the samples detected as saccades or blinks were removed from analysis (*n* = 286 out of 2440 total trials). For the remaining trials, blinks were linearly interpolated from 60 ms before to 160ms after the detected blinks. Saccades were linearly interpolated from 60 ms before to 60 ms after any detected saccade. Pupillary responses were then baseline normalized on a trial-by-trial basis to account for a downward drift in baseline that tends to occur across a task and for individual differences in pupil dynamic range (Winn et al., 2018). Baseline pupil size was defined as the average pupil size in the 1000 ms period prior to sound onset using the equation 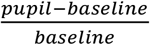.

Pupillary responses were then averaged across each SNR per participant, such that each subject had a single averaged pupillary response per SNR (9, 6, 3, and 0 dB). Pupillary responses from 0ms to 2480ms time-locked to the start of the speech stimulus were analyzed using growth curve analysis (GCA,(Mirman, 2017)). GCA uses orthogonal polynomial time terms to model distinct functional forms of the pupillary response over time. A GCA was fit using a second-order orthogonal polynomial to model the interaction with SNR level. This second-order model provides three parameters to explain the pupillary response. The first is the intercept, which refers to the overall change in the pupillary response over the time-window of interest. The second is the linear term (ot1), which represents the slope of the pupillary response over time. The third is the quadratic term (ot2), which represents the curvature of the pupillary response during listening.

The GCA model included fixed effects of SNR (reference = 0), Group (reference = YA), the interaction of SNR and Group, and all two- and three-way interactions between SNR, Group, and polynomial time terms. The random effect structure included a random intercept of subject on each time term and a random slope of the interaction between subject and SNR on each time term. GCA were conducted in R v4.2.2 (R Core Team, 2022) using the *lme4* package v1.1-31 (Bates et al., 2015) and *p*-values were estimated using the *lmerTest* package v3.1-3 (Kuznetsova et al., 2017).

(Pupil ~ (ot1 + ot2)*SNR*Group) + (ot1 + ot2|Sub) + (ot1 + ot2|Sub:SNR)

### Statistical Analyses

All analyses were conducted in R v4.2.2.

#### Group comparisons

Normality was evaluated using Shapiro–Wilk tests and Q–Q plots; homogeneity of variance was tested with Levene’s test. When sphericity assumptions were violated, Greenhouse–Geisser corrections were applied. Repeated-measures ANOVAs were used for within-subject factors (e.g., SNR, modulation depth), with false-discovery rate correction for multiple comparisons. Nonparametric tests (Mann–Whitney U) were used when normality assumptions were violated.

#### Regression models

Fifty-four participants (27 young adults, 27 middle-aged) were used in model building to ensure a balanced dataset across all metrics. To predict pupil dilation (3 dB SNR), multiple univariate models were built using pure-tone average (≤4 kHz), extended high-frequency thresholds (8–16 kHz), FMFR power, and FMFR discriminability as predictors, and the R^2^ were calculated. Predictors were then entered into a multivariate model iteratively based on highest univariate R^2^, and the adjusted R^2^ was calculated. Variables were z-scaled, and model performance was evaluated using adjusted R^2^ and likelihood-ratio tests. All major assumptions were examined, including normality of residuals, linearity, homogeneity of variance, and multicollinearity. Outliers were identified via Tukey’s fences (1.5×IQR) but retained when values were physiologically plausible.

#### Principal component reduction

For regression analyses, FMFR power and neural discriminability values across FM depths were reduced via principal component analysis (PCA). All variables were z-scored. Missing values were imputed using regularized iterative PCA(Josse and Husson, 2012a), with the number of components selected using generalized cross-validation(Bro et al., 2008; Josse and Husson, 2012b). Imputation quality was monitored via mean absolute error and root mean square error. We then ran unrotated PCA on the completed matrix and retained the first two components (eigenvalues > 1), which explained 35.9% and 29.3% of the variance, respectively. Loadings (between.702 and.843 on their respective components) and cosine squared (between.493 to.744 on their respective components) indicated that Dimension 1 captured FMFR power across modulation depths, whereas Dimension 2 captured FMFR neural discriminability. Accordingly, each participant’s FMFR data were summarized using these two scores for downstream analyses.

## Results

### Neural representations of temporal fine structure decrease with age despite normal hearing thresholds

Young (18–25 years, YA) and middle-aged (40–55 years, MA) adults (Fig. 1A) had clinically normal audiograms through 8 kHz, with mild-moderate threshold elevations at extended high frequencies (Fig. 1B, Table 1, 2). To quantify TFS encoding, we recorded FMFRs to a 520-Hz carrier modulated at 2 Hz with deviation depths from 0 (pure tone) to 20 Hz. FMFRs were analyzed using a spectrally specific frequency-demodulation approach (Parida et al., 2021) that isolates phase-locked neural power precisely along the known modulation trajectory (Fig. 1C). Two complementary indices of neural coding fidelity were derived. First, a power ratio metric compared response power along the FM trajectory to noise-floor power in frequency bins ±15 Hz away (Fig. 1D). Second, a neural discriminability index quantified how separable FM-evoked responses were from pure-tone responses, by comparing FM power fitted to the actual modulation trajectory versus an unmodulated pure tone of the same frequency. Neural discriminability was quantified by the average rate of change of a fitted sigmoid function of discriminability across depths (Fig. 1E–F).

**Table 1.**
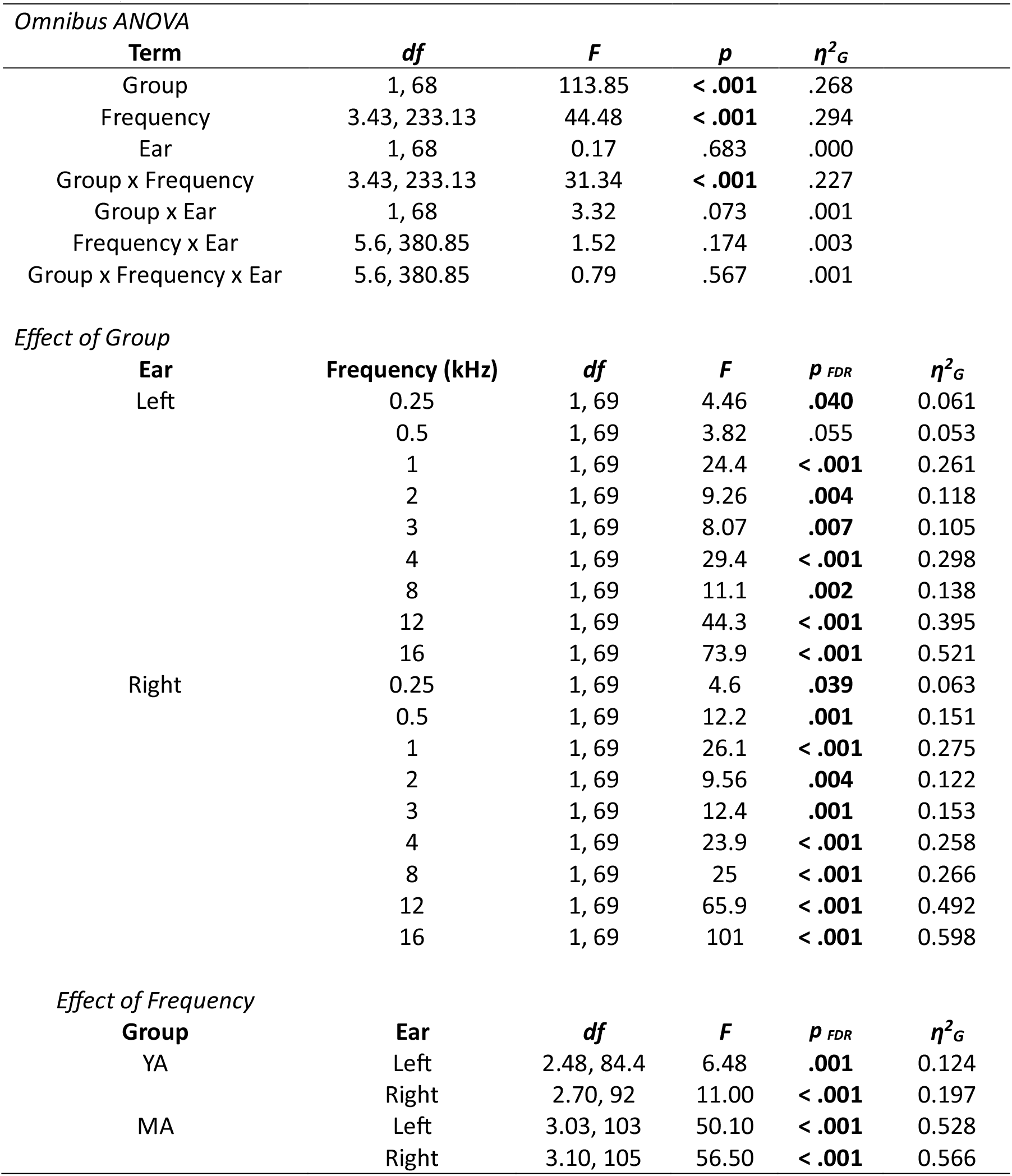
Comparison of air conduction thresholds using a three-way analysis of variance (ANOVA) (YA = 35, MA = 35)

**Table 2.**
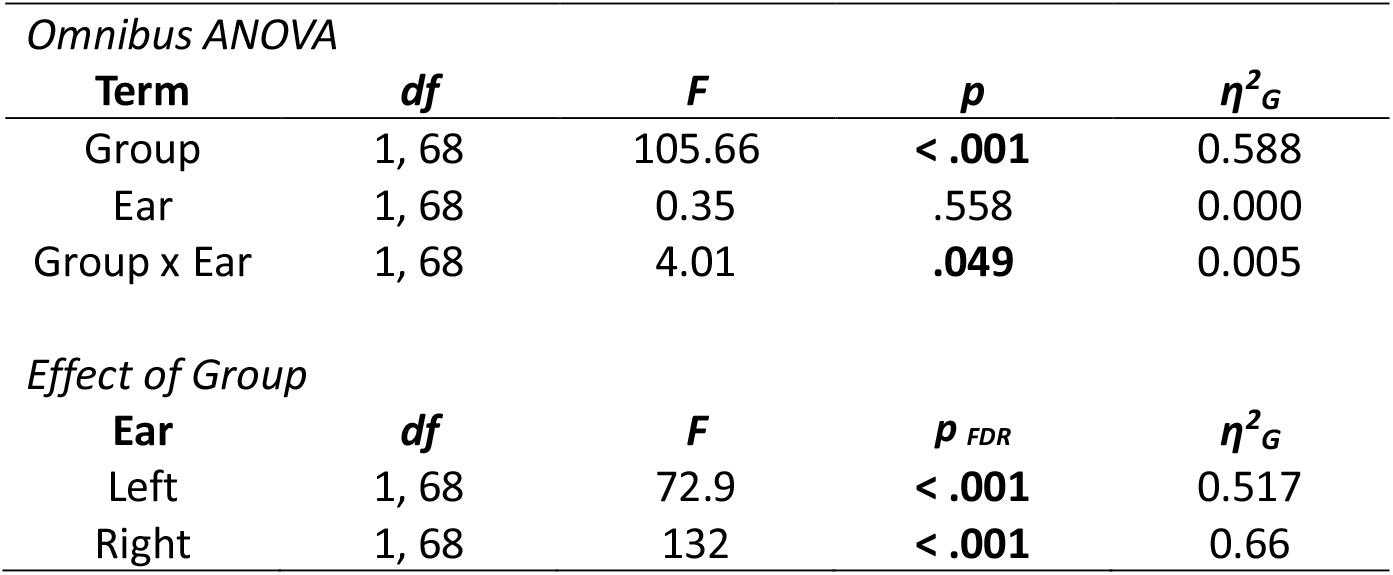
Comparison of extended high frequencies using two-way ANOVA (YA = 35, MA = 35)

| *Omnibus ANOVA* |  |  |  |  |
| --- | --- | --- | --- | --- |
| **Term** | ***df*** | ***F*** | ***p*** | **$\eta^2_G$** |
| Group | 1, 68 | 105.66 | **< .001** | 0.588 |
| Ear | 1, 68 | 0.35 | .558 | 0.000 |
| Group x Ear | 1, 68 | 4.01 | **.049** | 0.005 |
| *Effect of Group* |  |  |  |  |
| **Ear** | ***df*** | ***F*** | ***p<sub>FDR</sub>*** | **$\eta^2_G$** |
| Left | 1, 68 | 72.9 | **< .001** | 0.517 |
| Right | 1, 68 | 132 | **< .001** | 0.66 |

**Figure 1.**
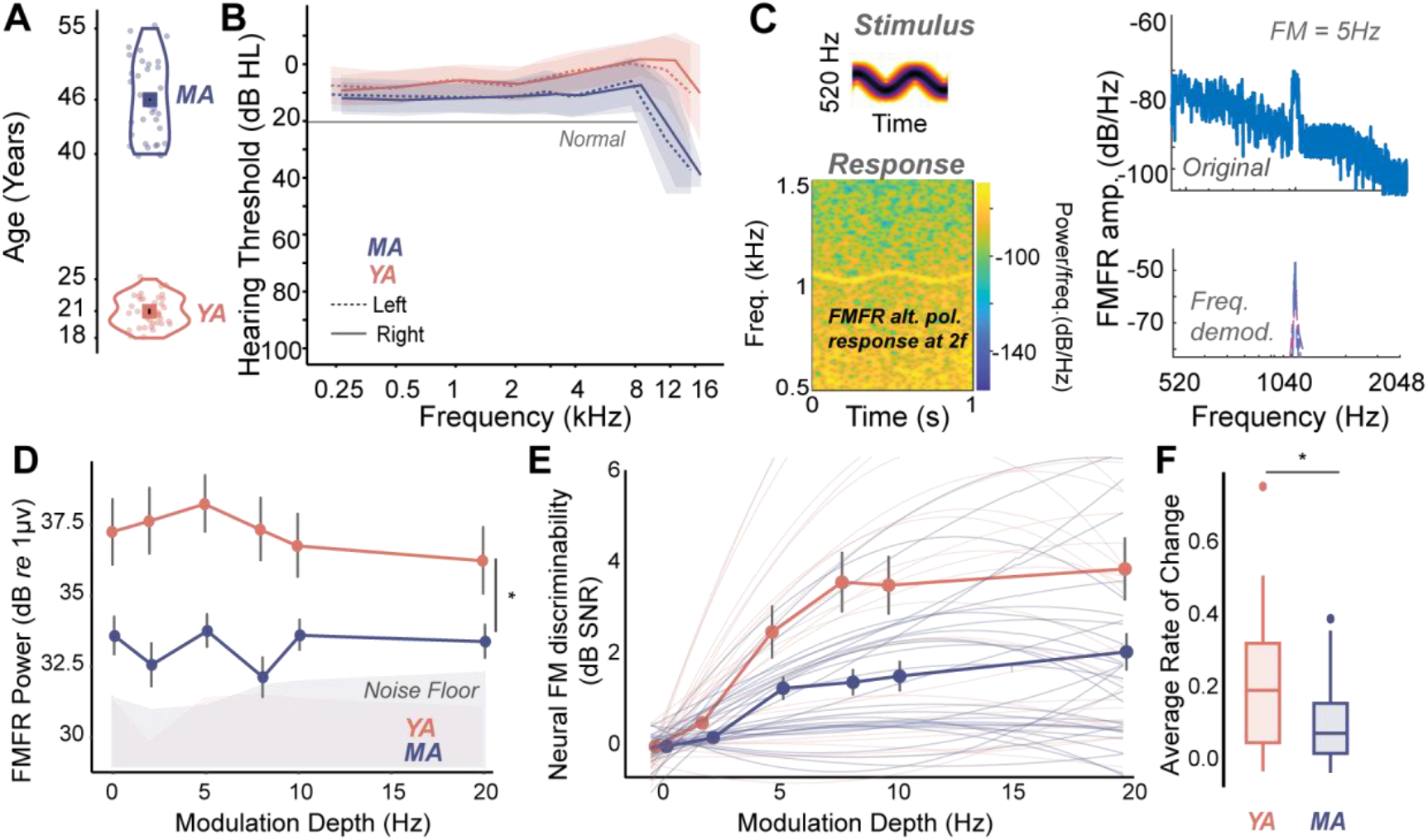
Neural representations of temporal fine structure decline with age despite normal hearing thresholds. (A) Participant age groups: young adults (YA, 18–25 yrs) and middle-aged adults (MA, 40–55 yrs). (B) Group-mean audiograms showing normal hearing through 8 kHz and mild threshold elevations at extended high frequencies in MA listeners. (C) Schematic of the spectrally specific frequency-demodulation method used to isolate frequency-modulation following responses (FMFRs) along the known modulation trajectory. (D) FM-locked neural phase-locking power (ratio of power along the FM trajectory vs. ±15 Hz flanking bins) significantly decreases with age. (E) Neural discriminability between FM and pure-tone responses. (F) Mean rate of change from the discriminability indices reveal significantly reduced phase-locking fidelity in MA listeners. Error bars = +SEM.

Both measures revealed systematic age-related declines in TFS encoding. MA listeners showed significantly reduced FM-locked power (Fig. 1D), with both a significant effect of age, and reduced FMFR power at all FM depth (p=0.001, ANOVA, Table 3). MA listeners also exhibited shallower discriminability slopes relative to YA listeners (Fig. 1E). There was a significant group effect on neural discriminability, and reduced discriminability at all FM depths (p = 0.007, ANOVA, Table 4). In addition, there was a significant reduction in the average rate of change of neural discriminability as well (t-test, YA = 27, MA = 30, t(42.367) = 2.386, p = 0.022).

**Table 3.** Comparison of FMFR power using two-way ANOVA (YA = 28, MA = 30)

| *Omnibus ANOVA* |  |  |  |  |
| --- | --- | --- | --- | --- |
| **Term** | ***df*** | ***F*** | ***p*** | **$\eta^2_G$** |
| Group | 1, 53 | 11.494 | **.001** | 0.143 |
| FM Depth | 5, 265 | 2.099 | .066 | 0.009 |
| Group x FM Depth | 5, 265 | 1.932 | .089 | 0.008 |
| *Effect of Group* |  |  |  |  |
| **FM Depth (Hz)** | ***df*** | ***F*** | ***p<sub>FDR</sub>*** | **$\eta^2_G$** |
| 0 | 1, 56 | 7.45 | **.012** | 0.117 |
| 2 | 1, 55 | 12.9 | **.001** | 0.19 |
| 5 | 1, 56 | 15.1 | **.001** | 0.212 |
| 8 | 1, 54 | 15.6 | **.001** | 0.224 |
| 10 | 1, 56 | 6.62 | **.016** | 0.106 |
| 20 | 1, 56 | 4.77 | **.033** | 0.078 |

**Table 4.** Comparison of Neural Discriminability using two-way ANOVA (YA = 25, MA = 26)

| *Omnibus ANOVA* |  |  |  |  |
| --- | --- | --- | --- | --- |
| **Term** | ***df*** | ***F*** | ***p*** | **$\eta^2_G$** |
| Group | 1, 49 | 7.801 | **.007** | 0.084 |
| FM Depth | 2.31, 113.25 | 34.713 | **< .001** | 0.230 |
| Group x FM Depth | 2.31, 113.25 | 4.729 | **.008** | 0.039 |
| *Effect of Group* |  |  |  |  |
| **FM Depth (Hz)** | ***df*** | ***F*** | ***p<sub>FDR</sub>*** | **$\eta^2_G$** |
| 2 | 1, 52 | 6.36 | **.025** | 0.109 |
| 5 | 1, 55 | 4.07 | **.049** | 0.069 |
| 8 | 1, 52 | 9.65 | **.015** | 0.157 |
| 10 | 1, 55 | 7.98 | **.018** | 0.127 |
| 20 | 1, 55 | 5.36 | **.030** | 0.089 |
| *Effect of FM Depth* |  |  |  |  |
| **Group** | ***df*** | ***F*** | ***p<sub>FDR</sub>*** | **$\eta^2_G$** |
| YA | 2.01, 48.1 | 21.4 | **< .001** | 0.257 |
| MA | 5, 125 | 13.7 | **< .001** | 0.231 |

Thus, there was a marked deterioration in neural phase-locking to frequency modulation by mid-life, reflecting degraded auditory-nerve and early brainstem synchrony, even when hearing thresholds remain clinically “normal”.

### Listening effort increases with age despite matched behavioral performance

To assess speech perception under multi-talker conditions, we used a four-digit comprehension task (Fig. 2A) which emphasizes reliance on TFS cues for speaker segregation (Parthasarathy et al., 2020). Participants tracked a target voice (fundamental frequency or F_0_ = 115Hz) while ignoring two simultaneous, co-localized distractors with flanking F_0s_ (175 Hz and 90 Hz). Signal-to-noise ratios (SNRs) were chosen such that overall accuracy declined with increasing difficulty, but performance was matched across groups (Fig. 2B, Table 5). This allowed us to isolate age-related changes in effort independent of behavioral performance.

**Table 5.**
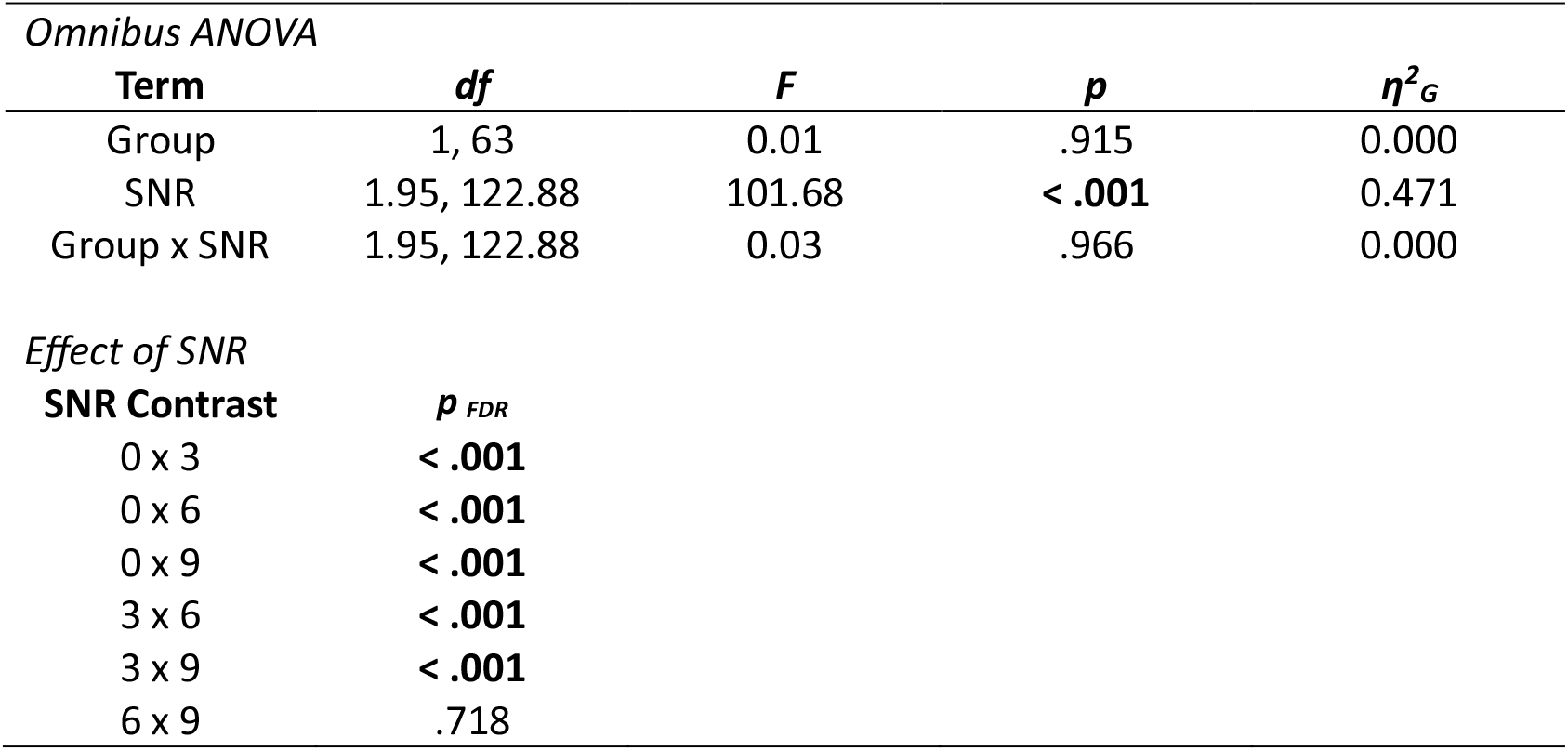
Comparison of multi-talker speech intelligibility using two-way ANOVA (YA = 31, MA = 34)

**Figure 2.**
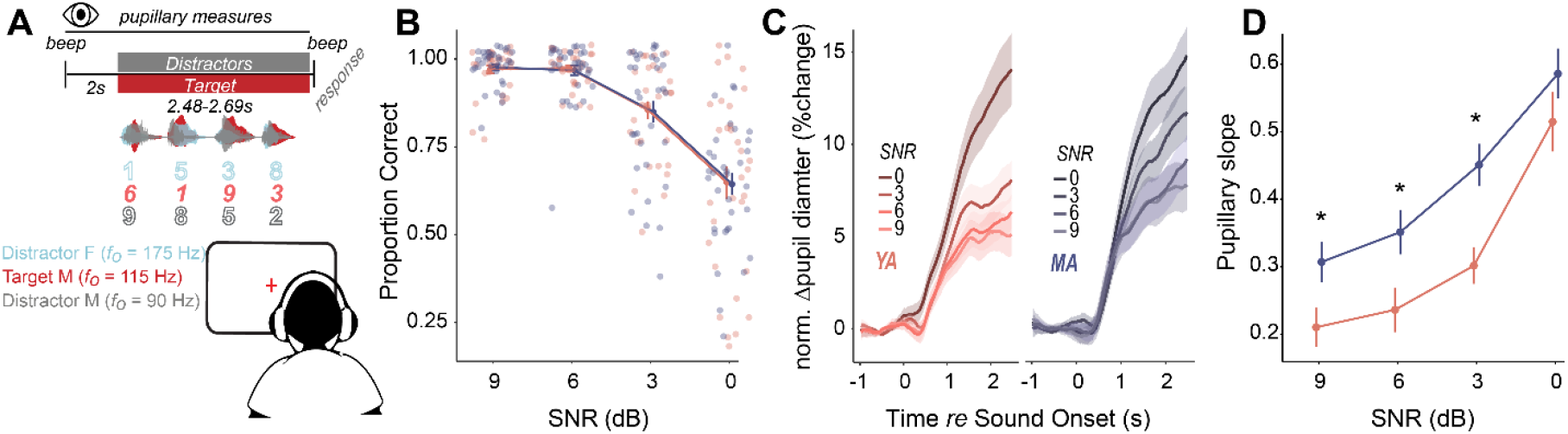
Listening effort increases with age despite matched behavioral performance. (A) Multi-talker four-digit comprehension task: participants report digits from a target male talker while ignoring two co-localized distractors differing in fundamental frequency (F_0_). (B) Speech intelligibility as a function of signal-to-noise ratio (SNR). Accuracy declines with increasing difficulty but remains statistically matched across age groups. (C) Mean pupil-dilation time courses normalized to pre-trial baseline, showing robust increases with task difficulty across listeners. (D) Pupil dilation slopes as a function of SNR. Middle-aged adults exhibit significantly larger dilations even when performance is near ceiling, indicating elevated listening effort independent of behavioral success.

Task-evoked pupil dilation, normalized to pre-trial baseline, served as an objective index of listening effort. Pupil diameter increased during listening with task difficulty across both groups, confirming sensitivity of the measure to cognitive load (Fig. 2C). However, MA listeners exhibited substantially larger dilations during listening than YA listeners even at difficulty levels where accuracy approached ceiling, as indexed by task-linked changes in pupillary slopes extracted using a growth curve analysis (Fig. 2D, Table 6, 7). Larger pupil-indexed effort was observed in SNRs 9, 6 and 3 dB in MA listeners. Pupillary slopes increased and were comparable in both YA and MA listeners at the hardest difficulty level of SNR 0.

**Table 6.** Comparison of pupillary slopes as a function of SNR and age group using a 2-way ANOVA (YA = 31, MA =34)

| *Omnibus ANOVA* |  |  |  |  |
| --- | --- | --- | --- | --- |
| **Term** | **df** | **F** | **p** | **$\eta^2_G$** |
| Group | 1, 59 | 57.50 | < .001 | 0.07 |
| SNR | 2.6, 153.11 | 24.59 | < .001 | 0.28 |
| Group X SNR | 2.6, 153.11 | 0.50 | .659 | 0.01 |
| *Effect of Group* |  |  |  |  |
| **SNR** | **df** | **F** | **$p_{FDR}$** | **$\eta^2_G$** |
| 9 | 1, 62 | 5.23 | .035 | 0.078 |
| 6 | 1, 59 | 6.26 | .030 | 0.096 |
| 3 | 1, 63 | 13.1 | .002 | 0.172 |
| 0 | 1, 63 | 1.55 | .218 | 0.024 |
| *Effect of SNR* |  |  |  |  |
| **Group** | **df** | **F** | **$p_{FDR}$** | **$\eta^2_G$** |
| YA | 2.26, 65.5 | 13.1 | < .001 | 0.297 |
| MA | 3, 90 | 11.9 | < .001 | 0.263 |
| *SNR contrasts* |  |  |  |  |
| **Group** | **SNR Contrast** | **$p_{FDR}$** |  |  |
| YA | 9 x 6 | .590 |  |  |
|  | 9 x 3 | .085 |  |  |
|  | 6 x 3 | .209 |  |  |
|  | 9 x 0 | < .001 |  |  |
|  | 6 x 0 | < .001 |  |  |
|  | 3 x 0 | < .001 |  |  |
| MA | 9 x 6 | .352 |  |  |
|  | 9 x 3 | .005 |  |  |
|  | 6 x 3 | .044 |  |  |
|  | 9 x 0 | < .001 |  |  |
|  | 6 x 0 | < .001 |  |  |
|  | 3 x 0 | .006 |  |  |

**Table 7.** Fixed effect estimates from the growth curve model of pupillary responses (YA = 31, MA = 34)

| **Fixed Effect** | ***b*** | ***SE*** | ***t*** | ***df*** | ***p*** | **95% CI** |
| --- | --- | --- | --- | --- | --- | --- |
| Intercept | 0.07 | 0.01 | 8.87 | 137.31 | < .001 | [0.06, 0.09] |
| ot1 | 0.51 | 0.06 | 9.34 | 133.67 | < .001 | [0.41, 0.62] |
| ot2 | -0.03 | 0.02 | -1.09 | 187.58 | .276 | [-0.07, 0.02] |
| SNR 3 | -0.03 | 0.01 | -3.70 | 189.58 | < .001 | [-0.05, -0.01] |
| SNR 6 | -0.04 | 0.01 | -4.91 | 190.09 | < .001 | [-0.06, -0.02] |
| SNR 9 | -0.04 | 0.01 | -5.24 | 189.58 | < .001 | [-0.06, -0.03] |
| Group (YA vs. MA) | 0.00 | 0.01 | 0.17 | 137.31 | .868 | [-0.02, 0.02] |
| ot1 x SNR 3 | -0.21 | 0.05 | -4.10 | 189.34 | < .001 | [-0.32, -0.11] |
| ot1 x SNR 6 | -0.28 | 0.05 | -5.33 | 189.80 | < .001 | [-0.38, -0.18] |
| ot1 x SNR 9 | -0.30 | 0.05 | -5.85 | 189.34 | < .001 | [-0.41, -0.2] |
| ot2 x SNR 3 | -0.03 | 0.03 | -1.08 | 190.05 | .284 | [-0.08, 0.02] |
| ot2 x SNR 6 | -0.03 | 0.03 | -0.95 | 190.83 | .344 | [-0.08, 0.03] |
| ot2 x SNR 9 | -0.03 | 0.03 | -1.21 | 190.05 | .226 | [-0.08, 0.02] |
| ot1 x Group | 0.07 | 0.08 | 0.93 | 133.67 | .352 | [-0.08, 0.22] |
| ot2 x Group | -0.05 | 0.03 | -1.65 | 187.58 | .101 | [-0.12, 0.01] |
| Group x SNR 3 | 0.01 | 0.01 | 1.32 | 189.58 | .187 | [-0.01, 0.04] |
| Group x SNR 6 | 0.01 | 0.01 | 1.22 | 190.70 | .224 | [-0.01, 0.04] |
| Group x SNR 9 | 0.01 | 0.01 | 1.09 | 189.97 | .276 | [-0.01, 0.03] |
| ot1 x Group x SNR 3 | 0.08 | 0.07 | 1.09 | 189.34 | .278 | [-0.06, 0.22] |
| ot1 x Group x SNR 6 | 0.05 | 0.07 | 0.62 | 190.36 | .535 | [-0.1, 0.19] |
| ot1 x Group x SNR 9 | 0.02 | 0.07 | 0.33 | 189.70 | .743 | [-0.12, 0.17] |
| ot2 x Group x SNR 3 | 0.06 | 0.04 | 1.75 | 190.05 | .081 | [-0.01, 0.14] |
| ot2 x Group x SNR 6 | 0.04 | 0.04 | 1.17 | 191.69 | .245 | [-0.03, 0.12] |
| ot2 x Group x SNR 9 | 0.02 | 0.04 | 0.61 | 190.57 | .546 | [-0.05, 0.09] |

These results reveal that middle-aged listeners recruit additional cognitive resources to maintain comparable comprehension in multi-talker scenarios, consistent with compensatory listening strategies in early auditory aging.

### Decreased neural coding of TFS cues predict increased listening effort during multi-talker listening

Finally, we tested whether neural encoding of temporal fine structure (TFS) predicted increases in listening effort. A multiple-regression model predicted pupil dilation at 3 dB SNR (the most challenging SNR with matched performance and increased listening effort in MA) using pure-tone average thresholds (PTA, ≤ 4 kHz), FMFR power, and FMFR discriminability. The model accounted for approximately 24% of the variance in pupil-indexed effort (MSE = 0.038, r^2^ = 0.249, p <0.001, Fig. 3A), indicating contributions from both subtle threshold elevations and neural synchrony deficits. To isolate the specific contribution of TFS encoding, we computed a residualized pupil measure controlling for PTA. Residual pupil diameter remained significantly correlated with FMFR power (Fig. 3B), linking weaker neural phase-locking directly to greater cognitive load during multi-talker listening. These results establish that degraded neural representations of TFS independently predict elevated listening effort, providing a mechanistic link between early auditory neural decline and everyday listening fatigue even in the absence of overt hearing loss.

**Figure 3.**
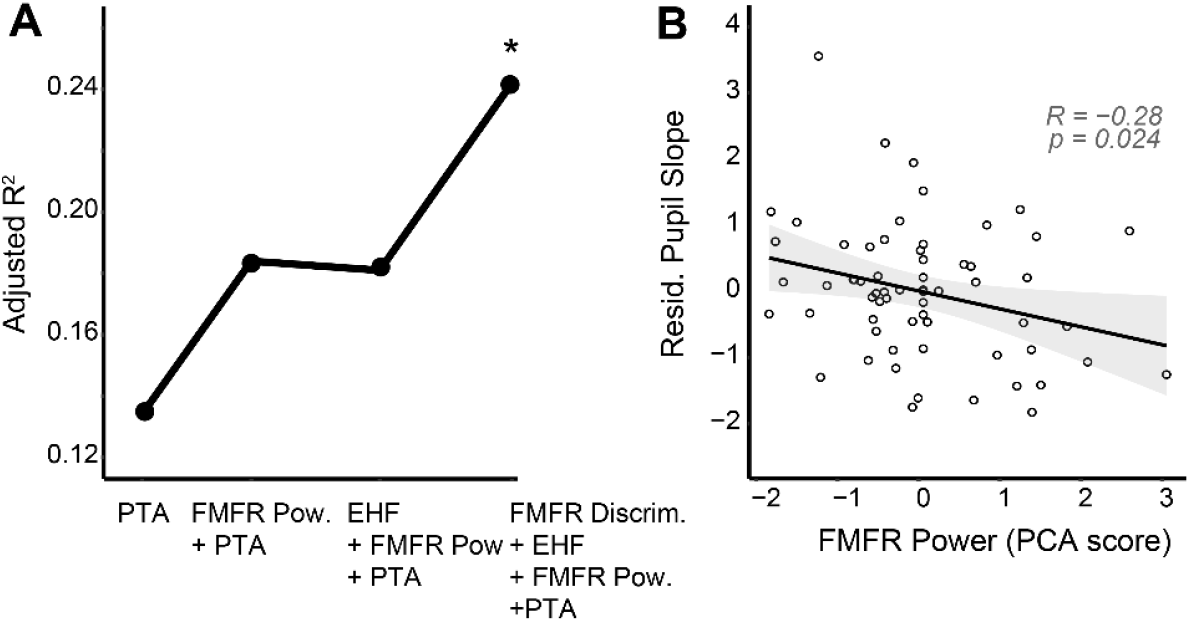
Decreased neural coding of TFS cues predict increased listening effort during multi-talker listenin. (A) Multiple-regression model predicting pupil dilation (3 dB SNR) from pure-tone thresholds ≤ 4 kHz, extended-high-frequency thresholds, FMFR power, and FMFR discriminability. The model explains ~24% of the variance in listening effort, implicating both subclinical threshold changes and neural TFS fidelity. (B) Residualized pupil diameter, corrected for audiometric factors, remains significantly correlated with FMFR power. Listeners with weaker neural phase-locking exhibit greater pupil-indexed effort, linking degraded peripheral temporal coding to increased cognitive load during multi-talker speech perception.

## Discussion

Here we demonstrate that neural representations of temporal fine structure (TFS) cues decline sharply by midlife, even in adults with clinically normal hearing thresholds. Using frequency-modulation following responses (FMFRs) to quantify neural phase-locking to low-frequency FM tones, we found that middle-aged listeners exhibited reduced FMFR amplitudes and shallower discriminability slopes compared to young adults, despite comparable audiometric profiles and speech-in-noise performance. Critically, these declines in neural synchrony predicted increased pupil-indexed listening effort during multi-talker speech perception, even when behavioral accuracy was matched across groups. Together, these findings identify a physiological correlate of subclinical auditory aging and establish a mechanistic link between degraded neural temporal coding and increased cognitive listening effort in everyday communication.

### Hidden hearing deficits and the gap in current diagnostics

A growing body of evidence suggests that age-related auditory decline begins well before measurable threshold loss on a standard audiogram (Hind et al., 2011; Tremblay et al., 2015; Spehar and Lichtenhan, 2018; Parthasarathy et al., 2020; Cancel et al., 2023). Many middle-aged adults report speech-in-noise difficulties that remain unexplained by conventional clinical tests such as pure-tone audiometry, emphasizing middle-age as a critical time window for early diagnostics. Further, behavioral speech-in-noise tests primarily capture performance accuracy rather than the effort required to achieve that performance, thereby underestimating the perceptual and cognitive burden imposed by subclinical neural degradation (McGarrigle et al., 2017; Peelle, 2018). Our data support this interpretation as well. Although middle-aged adults performed as well as younger listeners on a multi-talker comprehension task (Fig. 2B), they exhibited significantly larger task-evoked pupil dilations (Fig. 2D), reflecting greater recruitment of central resources to maintain comparable performance levels. These hidden physiological deficits masked by compensatory effort and resulting in a dissociation between performance and effort represent an under-recognized clinical challenge which leaves many individuals without a clear explanation or effective intervention for their listening difficulties.

### Temporal fine structure coding as a window into early neural decline

TFS cues play a fundamental role in parsing complex acoustic environments, providing fine-grained temporal information necessary for pitch perception, source segregation, and spatial localization (Moore, 2008; Oxenham, 2018). The ability of auditory nerve fibers to phase-lock to the fine structure of sound waves underlies this representation, and this neural synchrony may degrade with age, even when outer-hair-cell function remains intact (Joris et al., 2004; Ruggles et al., 2012; Bharadwaj et al., 2015; Borjigin and Bharadwaj, 2025). Behavioral assays such as low-frequency FM detection are primarily used to assess TFS sensitivity (Sek and Moore, 1995; Moore and Sek, 1996; Lorenzi et al., 2009). However, debate exists about whether these measures may be confounded by recovered envelope cues arising from cochlear filtering nonlinearities, making it difficult to isolate true neural phase-locking contributions (Verschooten et al., 2019). The FMFR overcomes this limitation by directly quantifying neural synchrony to FM tones using EEG, minimizing contamination from envelope cues because the analysis focuses on the carrier-related frequency region rather than the low-frequency modulation rate (Parthasarathy et al., 2020).

Our results extend previous work by demonstrating that even in listeners with clinically normal thresholds, the fidelity of FMFR phase-locking declines significantly by midlife. This decrease in phase-locking may be due to cochlear deafferentation, which can reduce the overall power of phase-locking at the level of the auditory nerve (Parthasarathy and Kujawa, 2018; Märcher-Rørsted et al., 2022; Ponsot et al., 2024; Temboury-Gutierrez et al., 2024). We used two measures to quantify the FMFR – overall power and neural discriminability (Fig. 1D-F). Middle-aged adults exhibited decreased response amplitudes in both metrics. The degree of loss with middle-age, especially in the neural discriminability index, was consistent across modulation depths. This suggests that the phase-locking ability may be decreased due to a loss of afferent connections, but the remaining auditory nerve fibers are likely intact in terms of their temporal processing ability. Single unit studies from animal models of cochlear deafferentation support this interpretation (Heeringa et al., 2020).

### Linking degraded TFS coding to listening effort

Our combined physiological and pupillometric findings provide a mechanistic bridge between subclinical neural degradation and the subjective experience of listening effort. The pupil response is a well-validated index of cognitive load, likely mediated by the locus-coeruleus– noradrenergic system (Aston-Jones and Cohen, 2005; Zekveld et al., 2010). Larger dilations in the absence of performance decrements therefore reflect compensatory recruitment of attentional and working-memory resources (Pichora-Fuller et al., 2016). We observed that listeners with weaker FMFR responses exhibited greater pupil dilation, even after accounting for small individual differences in audiometric thresholds (Fig. 3B), suggesting that degraded neural phase-locking to TFS cues directly increases cognitive demand during speech perception. This relationship parallels earlier reports showing that spectral or temporal degradation of speech elicits larger pupil responses (Zekveld et al., 2011; Zekveld and Kramer, 2014; Winn et al., 2015) and supports the hypothesis that impaired temporal coding forces greater reliance on top-down mechanisms to maintain comprehension (Shinn-Cunningham and Best, 2008; Peelle, 2018; Borjigin and Bharadwaj, 2025; Bramhall et al., 2025).

Regression analyses further indicated that subtle audiometric changes and reduced FMFR strength each explained unique variance in listening effort, together accounting for roughly one-quarter of the observed variability (Fig. 3A). Extended high-frequency thresholds, by contrast, did not contribute additional explanatory power once pure-tone averages were controlled. We have previously shown similar results in a middle-aged cohort, where PTA explained variance in speech in noise performance, but extended high frequencies did not, once those subtle changes in lower frequency thresholds are accounted for (Zink et al., 2024). The persistence of a robust correlation between FMFR metrics and pupil dilation after residualizing out audiometric effects underscores that TFS degradation represents an independent neural factor underlying hidden listening effort.

### Clinical implications and future directions

Auditory perception depends not only on peripheral fidelity but also on dynamic allocation of cognitive resources, particularly in noisy environments that require selective attention and working-memory updating (Gordon-Salant and Cole, 2016; Pichora-Fuller et al., 2016). The increased pupil-indexed effort we observed in middle-aged adults suggests that even modest sensory degradation can impose measurable cognitive load, potentially accelerating fatigue or disengagement from social communication. Chronic increases in listening effort may, in turn, contribute to reduced auditory stimulation and downstream cognitive consequences, linking hearing loss to dementia (Livingston et al., 2020). By identifying neural TFS degradation as an upstream contributor to this effort, our results highlight a potential early neural mechanism through which auditory aging interfaces with cognitive decline.

Current audiological evaluations rely almost exclusively on threshold-based measures that fail to detect temporal coding deficits. Even advanced clinical speech-in-noise tests (e.g., QuickSIN, WIN) primarily assess performance accuracy rather than effort, potentially missing subclinical neural impairments that listeners compensate for behaviorally. Integrating physiological measures such as the FMFR into the diagnostic workflow could enable earlier identification of individuals at risk for hearing-related cognitive decline. Moreover, quantifying listening effort through pupillometry or similar metrics may help evaluate the efficacy of hearing aids and auditory training programs that target temporal coding enhancement (Anderson et al., 2013; Bharadwaj et al., 2015).

Future work should directly link FMFR reductions to specific neural pathology such as cochlear deafferentation in animal models (Lopez-Poveda, 2014; Kujawa and Liberman, 2015) and identify additional cortical factors, both sensory - such as linguistic representations in the auditory cortex (Guo et al., 2025; McHaney et al., 2025), and suprasensory - such as attentional control or working-memory (Shinn-Cunningham and Best, 2008; Gordon-Salant and Cole, 2016), that can explain further unique contributions to increased listening effort. Together, these results position the FMFR as a sensitive, objective biomarker of early auditory neural decline, bridging peripheral sensory representations and the increased cognitive demands of everyday listening in midlife.

## Conflict of Interest

The authors declare no competing financial interests.

## Acknowledgements

This work was supported by the National Institute on Deafness and Other Communication Disorders-National Institutes of Health Grant R21DC018882 to A.P, Grant T32DC011499 to M.E.Z., the PNC-Trees Charitable Trust (PNC to B.C. and A.P.), and the National Institutes of Health (Grant F31DC020085-01A1 to J.R.M.). We thank Claire Mitchell, Olivia Flemm, Megan Hallihan, Kathryn Bergstrom, Sarah Anthony, and Shaina Wasileski for their assistance with participant recruitment and data collection, Jennifer Klara for administrative support, and the Clinical and Translational Science Institute at the University of Pittsburgh, supported by the NIH Clinical and Translational Science Award (CTSA) program, grant UL1 TR001857 for assistance with participant recruitment.

## Author contributions

LZ – data collection, analysis, stats, writing; SP – FMFR analysis; JM – data collection, methodology pupillometry analysis; MZ – data collection; BC – conceptualization, editing; AP – conceptualization, writing

## Data availability

Data and code used in this manuscript are available in Open Science Framework - https://osf.io/pca3x/overview

## Notes

### Competing Interest Statement

The authors have declared no competing interest.

